# Improving protein alignment algorithms using amino-acid hydrophobicities - Applications of TMATCH, A new algorithms

**DOI:** 10.1101/2020.01.01.878769

**Authors:** David P. Cavanaugh, Krishnan K. Chittur

## Abstract

**Motivation:** Sequence database search and matching algorithms are an important tool when trying to understand the structure (and so the function) of proteins. Proteins with similar structure and function often have very similar primary structure. There are however many cases where proteins with similar structure have very different primary structures. Substitution matrices (PAM, BLOSUM, Gonnett) can be used to identify proteins of similar structure, but they fail when the sequence similarity falls below about 25%.

**Results:** We have described a new algorithm for examining the the primary structure of proteins against a database of known proteins with a new hydrophobicity index. In this paper, we examine the ability of TMATCH to identify proteins of similar structure using sequence matching with the hydrophobicity index. We compare results from TMATCH with those obtained using FASTA and PSI-BLAST. We show that by using similarity patterns spread across the entire length of two proteins we get a more robust indicator of remote relatedness than relying upon high similarity scoring pair regions.

**Availability:** The program TMATCH is available on request

## 1 INTRODUCTION

Considerable time and effort has been expended to find and refine computer algorithms for the detection of proteins related to or homologous with a given search protein. These efforts have been expanded to facilitate searches of protein databases for such applications as phylogenetic analysis, protein classification, elucidation of cellular biochemical pathways by cross species comparative analysis and the illumination of unknown protein structure as inferred from homologous proteins of known structure.

The understanding of protein structure is a key step in understanding the function of proteins. Consequently, performing alignment algorithm based searching of protein databases is one of the key tools for theoretical biology, bio-technology and bio-medical research. Our TMATCH algorithm is an attempt to further the state of the art in this important field of finding related / homologous proteins.

TMATCH is founded upon a dynamic programming algorithm much like the Smith-Waterman algorithm and the popular database alignment search algorithms like FASTA. The TMATCH algorithm was specifically formulated to tackle the problem of finding remote protein homologues at or below the twilight zone (less than 25% sequence identity) where reliable detection of these remote homologues becomes very difficult. The TMATCH alignment algorithm leverages the extra amino-acid similarity information found within the hydrophobicity scale introduced in our previous TMATCH hydrophobicity paper (Cavanaugh 2015). This hydrophobicity scale provides a unified statistical and conceptual framework for protein alignment based on molecular geometry of the different amino-acid residues, their interaction with water and structural components of the organization of water around the folded proteins.

## 2 METHODS

### 2.1 Approach

Within this paper the emphasis is upon illustrating the performance features of the TMATCH algorithm. The details of the TMATCH algorithm and of the cognate hydrophobicity proclivity scale has been described in separate papers (Cavanaugh 2012b, Cavanaugh 2015). The performance of the TMATCH alignment algorithm is benchmarked / tested against several extant alignment methods, including database search methods (Smith-Waterman, BLAST and PSI-BLAST), with particular emphasis paid to the statistical significance / expectation functions. Two protein data sets were assembled as the databases against which the TMATCH searches were conducted. The first dataset is from a family of Tryptophan like Serine proteases curated by Pearson (Pearson 1997), which represent a particularly challenging dataset (because of the family primary sequence diversity) to benchmark TMATCH against the FASTA and Smith-Waterman algorithms (Pearson 1997 characterization).

Two of the proteins in the Tryptophan like Serine protease dataset were selected as the “search sequences” to generate the table of performance statistics. The second dataset was harvested from the PDB (PDB, Protein Databank) with a simple keyword search for “DNA polymerase.” A DNA polymerase B from a hyper-thermophilic, deep sea archaeon (*Thermococcus thioreducens*) was used as the search sequence and a table of performance statistics was created. The *T. thioreducens* PCNA protein was used as a search sequence in BLAST to obtain the HSSP alignments for a psychrophilic (cold loving) archaeon, *Methanoccoides burtonii*, and to see if the *Shigella boydii* bacteria Beta clamp protein would be detected using BLAST. The *T. thioreducens* PCNA protein search was repeated with PSI-BLAST with five iterations. The T. thioreducens PCNA protein BLAST / PSI-BLAST results were compared with TMATCH alignments of the *T. thioreducens* PCNA “search” protein against the *M. burtonii*, PCNA protein and the *S. boydii* bacteria Beta clamp protein.

### 2.2 TMATCH matricies

The TMATCH algorithm uses three matrices to support its implementation of the basic dynamic programming algorithm. These matrices are the match matrix, the cumulative score matrix and the backtrack matrix. The match matrix is the first step comparison of the amino-acids in the search (row) protein and the match (column) protein that is aligned against the search protein. The match, or residue to residue comparison, derives from a rescaled absolute value of delta (by subtraction) of hydrophobicities between each row / column cell intersection amino-acid pair. Each cell in the cumulative sum matrix represents the local optimal cumulative score representing the best path or trace from the beginning (upper left corner) of the cumulative sum matrix to that particular cell. Each cell’s value is determined by how that cell is entered, from the cell on the left, the cell above or the cell on the upper left. Entering the current cumulative score matrix cell from the left or above results in a gap opening penalty being assessed or added to the score at the previous cell.

If the current cumulative matrix cell is entered from the top left either a reward score (for favorable transition) or a penalty score (unfavorable transition) will be assessed by adding the reward / penalty to the previous cell’s score. The corresponding match matrix cell values are part of the calculation of the current cumulative score matrix cell value. The path chosen into the current cumulative matrix cell depends upon which transition / path results in the highest score for the current score matrix cell. Each cell of the back track matrix corresponds to each cell of the cumulative score matrix. The back track matrix cell values represent the direction the corresponding cumulative score matrix cells were entered to yield the best path transition having the highest score: −1 = enter from the cell to the left, 0 = enter the cell from the upper-left or +1 = enter from the cell above the current cell. In order to improve large protein database search times, TMATCH provides for a calibrated estimate of the %hydrophobic fuzzy match derived from the highest cumulative score in the cumulative score matrix, which eliminates the over-head of computing the back track matrix, extracting the alignment and calculating the cumulative %hydrophobic match derived from the pairwise comparison of each corresponding residue in the two aligned proteins. The estimate of the %hydrophobic fuzzy match is shown within the previous TMATCH theory paper (Cavanaugh 2012b) to be a robust and efficient estimate of the %hydrophobic match calculated from the extracted alignment.

### 2.3 TMATCH gap penalty, significance function and treatment of HSSP sequences in a dot plot like fashion

The FASTA and BLAST algorithms are devised to find, nucleate and grow aligned regions of high similarity, termed high scoring segment pairs, between the search protein and each incoming candidate match protein. These high scoring segment pair regions detect local regions of high similarity, such as would be introduced by regions of similar secondary structure. Because these regions of high similarity have enhanced visibility with the amino-acid hydrophobicity metric, they become quite visible as similarity patterns in diagonals within the match matrix analogous to patterns detected with dot plot analysis. The TMATCH algorithm takes advantage of the high similarity diagonal patterns by introducing a reward to the cell transition score to cause the path trace in the cumulative score and backtrack matrices to preferentially take these high similarity diagonal patterns.

The TMATCH algorithm uses a fixed gap penalty rather than an affine gap penalty as does FASTA and BLAST. The TMATCH algorithm uses an unfavorable diagonal transition penalty to outweigh the fixed gap penalty and force the opening of a gap. Unlike the FASTA and BLAST algorithms, TMATCH can detect similarity patterns across the entire length of two proteins being aligned, thereby capturing more information about the entire 3D fold, or basic scaffold, of the two proteins being aligned and compared than would be found by depending upon regions of high similarity alone. To the extent that TMATCH looks at similarity across the whole of two proteins being aligned with a metric containing information sensitive to secondary and tertiary structure, TMATCH will provide results which compare favorably with structural alignments. We demonstrate this in this paper with the results of the Tryptophan like Serine Proteases.

The alignment significance function F(Zb) has four defined regions of relationship by increasing degree of relatedness between the the search sequence and the candidate match sequences as explained below and in table 1. These definitions are being introduced to provide for meaningful interpretations of the statistical significance score. The interpretation of the ranges in table 1 is that the upper bound is specified and the lower bound is greater than the next table entry below it.

**Table 1.**
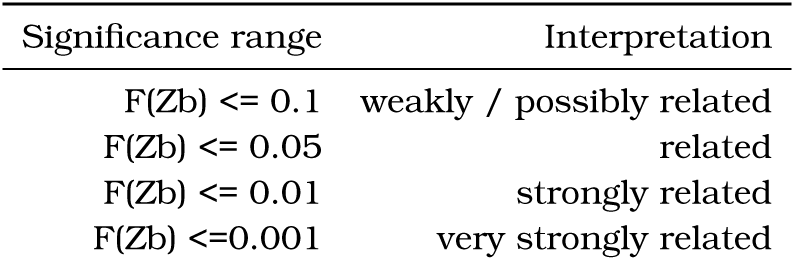
Interpretation of alignment statistical significance F(Zb)

| Significance range | Interpretation |
| --- | --- |
| $F(Zb) \leq 0.1$ | weakly / possibly related |
| $F(Zb) \leq 0.05$ | related |
| $F(Zb) \leq 0.01$ | strongly related |
| $F(Zb) \leq 0.001$ | very strongly related |

## 3 RESULTS

### 3.1 Tryptophan like Serine Proteases

The Trypsin-like Serine Protease super-family (table 3) is a very diverse group (at the level of their primary sequences) of proteins sharing the characteristic catalytic triad (D-H-S) and homologous 3D structures. A number of intra-super-family comparisons result in percent sequence matches around the “twilight zone” (threshold of 25% sequence similarity). It can thus be difficult to recover all 160 members of this super-family from a smaller set without using some advanced searching techniques. Since the tryptophan like serine proteases share the same basic fold / tertiary structure, this super-family poses an excellent test of the properties and desirable features of the TMATCH alignment algorithm. This set of proteins was carefully curated and the alignments characterized by Pearson (Pearson 1997) and those results serve as the basis of comparison to the TMATCH results in this section. The percent identities discussed in the text are from the percent identities reported in Table 3 are from the TMATCH alignment and are very similar to the percent identities in Pearson (Pearson 1997).

**Table 2.** Select sequence comparisons show an illustration of how a diverse protein family / super-family may need more than one search sequence and search pass

| Search Sequence -><br>(Matched sequence) | TRYP_BOVIN | FA7_RABBIT |
| --- | --- | --- |
| G156_PARPR | not significant | not significant |
| TRY1_ANOGA | significant | significant |
| PRLA_LYSEN | significant | significant |
| LORI_MOUSE | not significant | weakly significant |
| PLMN_PIG | not significant | significant |

**Table 3.** Relationships between TMATCH %Identity, %Hs, %%Hr and f(Zb) significances from two search sequences TRYP_BOVIN and FA7_RABBIT using Trypsin-like serine proteases (35 curated sequences from Pearson 1997). LORI_MOUSE and G156_PARPR are not serine-proteases. Compare the TRYP_BOVIN search sequence local alignment scores (E() same score algorithm as BLAST) and the F(Zb) significance levels in the third to the last and the last column. We see that the relative significance assessment between F(Zb) and the alignment score are for the most part similar in significance/interperation & meaning, except where there are differences reflected by the global alignment versus the regions of high similarity local alignments. Notice how the F(Zb) significance levels vary between TRYP_BOVIN and FA7_RABBIT as the respective search sequences. The F(Zb) significance levels are similar in some cases and in other cases the F(Zb) significance levels vary widely, which hammers home the point that some protein families, like the Tryptophan like Serine Proteases, are widely divergent in sequence identity and detecting all of the members of a family can be difficult, thus may take two or more search sequeqnces to recover the whole protein family.

| Protein | TRYP_BOVIN<br>%Hs match | TRYP_BOVIN<br>%Hr match | TRYP_BOVIN<br>%identity | TRYP_BOVIN<br>F(Zb) | FA7_RABBIT<br>F(Zb) | Pearson<br>TRYP_BOVIN E() |
| --- | --- | --- | --- | --- | --- | --- |
| TRY2_HUMAN | 80.38 | 78.33 | 70.42 | 0.00% | 65.68% | 0 |
| TRYP_PLEPL | 70.14 | 58.75 | 40 | 0.00% | 40.50% | 0 |
| KLK2_HUMAN | 64.26 | 60.42 | 39.58 | 0.01% | 52.24% | 0 |
| TRYP_FUSOX | 66.67 | 57.92 | 39.58 | 0.01% | 41.42% | 1.00E-27 |
| TRYM_CANFA | 63.11 | 59.58 | 35 | 0.01% | 7.37% | 1.00E-24 |
| TRYA_DROME | 65.38 | 56.67 | 39.58 | 0.01% | 6.79% | 1.00E-30 |
| RVVA_VIPRU | 65.94 | 55.42 | 32.92 | 0.01% | 75.10% | 0 |
| TRY1_ANOGA | 62.71 | 58.33 | 33.75 | 0.01% | 4.09% | 1.00E-31 |
| TRY5_ANOGA | 61.92 | 54.17 | 31.25 | 0.02% | 3.81% | 1.00E-27 |
| FA9_RAT | 59.81 | 56.25 | 38.33 | 0.02% | 3.31% | 1.00E-29 |
| TRYP_STRGR | 62.46 | 50.42 | 30.83 | 0.03% | 24.38% | 1.00E-19 |
| HGF_HUMAN | 62.46 | 50.42 | 30.83 | 0.03% | 24.38% | 1.00E-17 |
| CERC_SCHMA | 60.73 | 48.75 | 23.33 | 0.04% | 7.21% | 0.00001 |
| FA7_RABIT | 47.11 | 60.42 | 30.83 | 0.06% | --- | 1.00E-26 |
| PRTZ_BOVIN | 50.63 | 56.25 | 21.25 | 0.07% | 0.29% | 0.015 |
| PRTA_STRGR | 56.48 | 49.58 | 17.92 | 0.07% | 7.80% | 64 |
| ACH1_LONAC | 59.12 | 45.83 | 24.17 | 0.09% | 98.46% | 1.00E-15 |
| GSEP_BACLI | 53.06 | 51.25 | 21.67 | 0.10% | 1.62% | 0.45 |
| DLK_HUMAN | 48.64 | 53.33 | 21.25 | 0.15% | 0.45% | 9.9 |
| ACRO_PIG | 43.67 | 57.92 | 29.17 | 0.16% | 0.48% | 1.00E-25 |
| PRTB_STRGR | 53.8 | 46.67 | 17.5 | 0.19% | 3.07% | 16 |
| AGI_HORVU | 56 | 41.67 | 15.83 | 0.31% | 97.57% | 6.2 |
| PRLA_LYSEN | 48.28 | 49.17 | 20.83 | 0.32% | 0.30% | 3.1 |
| AGI_URTDI | 49.49 | 47.5 | 20.83 | 0.35% | 1.10% | 3.9 |
| URTB_DESRO | 44.94 | 51.25 | 16.67 | 0.41% | 0.04% | 1.00E-26 |
| PRTC_HUMAN | 40.84 | 50.83 | 17.08 | 0.98% | 0.01% | 1.00E-25 |
| KRUC_SHEEP | 55.08 | 33.75 | 15.83 | 1.75% | 100.00% | 1.3 |
| KCR8_YEAST | 31.28 | 53.33 | 19.17 | 4.35% | 0.19% | 4.7 |
| YB9X_YEAST | 23.56 | 59.58 | 24.17 | 6.04% | 0.40% | 9.5 |
| CFAB_MOUSE | 25.68 | 55.42 | 21.25 | 9.60% | 0.12% | 0.01 |
| AMY_CLOAB | 27.05 | 53.75 | 19.58 | 10.26% | 0.50% | 5.7 |
| CO2_HUMAN | 25.52 | 54.17 | 17.5 | 13.19% | 0.39% | 0.0001 |
| PQN25_CAEEL | 29.59 | 48.33 | 20.83 | 19.49% | 0.21% | 5.4 |
| PROS_MOUSE | 27.61 | 47.92 | 15.83 | 32.06% | 0.14% | 9.8 |
| PLMN_PIG | 22.97 | 52.5 | 20.42 | 32.39% | 0.66% | 1.00E-29 |
| LORI_MOUSE | 27.32 | 38.75 | 18.33 | 91.41% | 5.68% | 0.24 |
| G156_PARPR | 7.79 | 53.33 | 20.83 | 98.83% | 99.99% | 5 |

Advanced search techniques generally follow a search paradigm where protein A is statistically correlated (though alignment database searching) with protein B, protein B is statistically correlated with protein C in a similar fashion, therefore protein A is deemed to be statistically correlated with protein C by statistical transitivity. Thus, protein C is recovered through a search of a large search space by a two-step process with protein A as the first start search sequence and protein B as the second search sequence. Statistical transitivity relationships become unclear unless the percent similarity at each step of the statistical transitivity chain is very high. Moderately high statistical similarities in a hierarchical network of statistical transitivities roll off quickly obscuring the relationship between the protein at the root (the search sequence) and other members of the family just a few steps removed within the family tree. For example, in 4 steps of a statistical transitivity chain at an 85% statistical relationship in each step of the chain results in a 52% similarity between the first and fifth protein in the transitivity chain. With this multi-step type of database search paradigm, entire protein families / super-families can be recovered. Motif profiles can be derived once the family / super-family assemblage is complete, but these profiles may not be especially powerful for general database searches for other homologous proteins, where short profiles matches are statistically likely to be found owing to the sheer number of sequences occurring in the databases. (Pearson 1997)

In this tryptophan-like Serine protease super-family, search techniques can use two sequence regions: the Histidine active site is [LIVM]-[ST]-A-[STAG]-H-C and the Serine active site is G-D-S-G-G. These profiles are not long, so the problem of false alarms from searching many sequences exists for this super-family. However, within this Serine protease super-family homologous regions are generally spread throughout the proteins, rather than focused at the active site, a phenomenon expected for proteins that share homologous tertiary structure. Generally speaking, extended regions of lower level homology are more reliable ways of detecting homologous proteins than a few regions of locally, highly correlated subsequences (eg. sequences of high scoring segment pairs). (Pearson 1997)

After Pearson, we used thirty seven proteins with thirty five members in the tryptophan-like Serine protease super-family (Pearson 1997) for an alignment study with TMATCH, and then the results compared with that obtained using the Smith-Waterman algorithm (Pearson 1997). The Serine protease TRYP_BOVIN (bovine trypsinogen) was selected as the search sequence as it was in the original study (Pearson 1997). Twenty three of these Serine proteases were judged by BLAST to be significant matches at expectation values (< 0.02) The remaining 14 proteins were scored as marginally significant to non-significant (E()=0.24 -> 64.0). The least significant alignment match protein, PRTZ_BOVIN (vitamin K-dependent protein Z, E()=0.015), when used as the search sequence recovers a number of other members of the tryptophan like serine proteases family through statistically significant alignments using BLAST. Of the 14 proteins with an expectation value of 0.24 and up, four are linked by statistically significant alignments (these sequences used as the search sequence) with members of the Tryptophan-like Serine protease family and / or their tertiary structures: GSEP_BACLI, PRLA_LYSEN, PRTA_STRGR, PRTB_STRGR. (Pearson 1997)

We expect that proteins with homologous tertiary structure, but low primary sequence identity, should have a high degree of residue physico-chemical similarity, such as residue hydrophobicity (Cavanaugh 2008a). *Therefore we expected that TMATCH alignments for protein members of a protein family / super-family should recover high percentage hydrophobicity (%H) matches, even when the percentage identities are low and the FASTA / BLAST significance score with the selected search protein are not statistically significant*.

The percentage identities stretch from 20-75%, with a few of the sequences selected through a BLAST alignment algorithm being statistically non-significant matches to the TRYP_BOVIN search sequence, owing to low percentage identity (approximately 25%), but known to be homologous proteins from advanced search methods and confirmation by tertiary structural conservation (Pearson, 1997). Additionally, there is a wide variation in sequence lengths within this assemblage of proteins, many being significantly longer than the search sequence, which poses a significance test of the TMATCH alignment statistics and their correlation with conventional alignment metrics like percentage identity. The results of the comparison of TMATCH and the BLAST alignment statistics are summarized in Table 3.

In table 3, there are many sequences whose statistical significance E() scores are > 0.2 and thus require some additional evidence to support their status as homologous to the search sequence TRYP_BOVIN. Expectation E() scores are derived from local alignment HSP statistical extreme value theory (Altschul 1997, Karlin 1990).

As mentioned above four protein sequences (known to be homologous to the search sequence by advanced search methods using BLAST described above lie in the questionable area of detected, but non-statistically significant sequences: GSEP_BACLI, PRLA_LYSEN, PRTA_STRGR, PRTB_STRGR. All four of these sequences only have a 17-22% match with the search sequence, putting these sequences below the twilight zone threshold. However, each of these four sequences has a percent hydrophobicity fuzzy match of around 50 %, illustrating the enhanced information available in the hydrophobicity proclivity score, and not surprisingly these four Proteins were recovered by the TMATCH alignment with a strong statistical significance.

We are aware that an improved version of BLAST (PSI-BLAST) does exist which serves to automate the multi-pass, interactive searching for remote protein homologues. However, the TMATCH algorithm recovers all of these as statistically significant matches to the search sequence directly as seen in table 5. Two of the sequences (LORI_MOUSE and G156_PARPR) are not homologous to the search sequence (Pearson 1997) and do not share the same tertiary structure as the search sequence and are not statistically significant using the f(Zb) fit to the binomial distribution (see Table 3). Furthermore, these two proteins are known not to be in the tryptophan like serine protease super-family (Pearson 1997). *The TMATCH analysis suggests then that in order to conserve structural homology, nature substitutes amino-acids sharing strong physico-chemical similarities in distantly related proteins (eg. low % identity)*.

**Table 4.** Search sequence: hyper-thermophilic archaeon *T. thioreducen*s and DNA polymerase B match sequences from the PDB. It’s no suprise that the alignment significance scores for some archaeons are very significant reflecting the high similarity with the *T. thioreducens* DNA polymerase B. What is suprising is that the human herpes virus DNA polymerase B is so closely related to the DNA polymerase B of the archaeon *T. thireducens*. Notice the rise in the Alpha significance values from very strongly related through related, possibly related and onto unrelated. A careful examination of the protein descriptions shows a very nice correlation between the *T. thioreducens* DNA polymerase B significance levels and the corresponding relatedness as reveiled by the protein descriptions. For example, there are 3 proteins (a mouse mitocondrial polymerase, a human DNA polymerase gamma and an E. coli non-polymerase protein) in the possibly related alpha significance zone between 5% and 10%

| PDB accession # | # residues | %Hs match | %Hr match | % Identity | F(Zb) | Description |
| --- | --- | --- | --- | --- | --- | --- |
| 1QHT | 775 | 96.43 | 95.1 | 91.61 | 0.00% | DNA POLYMERASE FROM THERMOCOCCUS SP. 9ON-7 ARCHAEON |
| 1TGO | 773 | 95.55 | 93.81 | 89.81 | 0.00% | THERMOSTABLE DNA POLYMERASE B THERMOCOCCUS GORGONARIUS |
| 1WNS | 774 | 95.18 | 93.42 | 89.03 | 0.00% | DNA polymerase B from hyperthermophilic archaeon pyrococcus kodakaraensis KOD1 |
| 1WN7 | 774 | 95.05 | 93.29 | 88.9 | 0.00% | archaeal family B DNA polymerase mutant Thermococcus kodakarensis |
| 1QQC | 773 | 94.79 | 92 | 86.45 | 0.00% | ARCHAEBACTERIAL DNA POLYMERASE D.TOK |
| 1D5A | 733 | 89.82 | 85.94 | 79.87 | 0.00% | ARCHAEBACTERIAL DNA POLYMERASE D.TOK. Desulfurococcus |
| 2JGU | 775 | 89.21 | 85.94 | 76.77 | 0.00% | Dna Polymerase Pfu Pyrococcus Furiosus |
| 1S5J | 847 | 60.78 | 53.42 | 22.84 | 0.02% | DNA Polymerase B1 from the archaeon Sulfolobus solfataricus |
| 2GV9 | 1193 | 51.73 | 57.42 | 22.58 | 0.05% | Herpes Simplex virus type 1 DNA polymerase |
| 2HPM | 1220 | 51.18 | 57.55 | 23.35 | 0.05% | Polymerase III alpha - e. coli |
| 2DTU | 896 | 58.07 | 50.58 | 19.87 | 0.05% | proofreading beta hairpin loop found in exonuclease domain of DNA polymerase B family |
| 1Q8I | 783 | 58.54 | 49.16 | 19.35 | 0.06% | ESCHERICHIA coli DNA Polymerase II |
| 1IH7 | 903 | 57.91 | 49.68 | 18.45 | 0.06% | Apo bacteriophage RB69 DNA Polymerase |
| 1Q9Y | 906 | 58.02 | 49.55 | 18.97 | 0.06% | ENTEROBACTERIA PHAGE RB69 GP43 DNA POLYMERASE |
| 2P5G | 903 | 57.87 | 49.55 | 18.58 | 0.06% | DNA POLYMERASE FROM BACTERIOPHAGE RB69 gp43 |
| 1TAQ | 832 | 55.94 | 48.52 | 16.13 | 0.10% | STRUCTURE OF TAQ DNA POLYMERASE Thermus aquaticus |
| 1TAU | 832 | 55.99 | 48.13 | 15.61 | 0.10% | TAQ POLYMERASE Thermus aquaticus |
| 2HQA | 917 | 54.74 | 48.52 | 18.45 | 0.12% | catalytic alpha subunit of E. Coli - DNA polymerase III e. coli |
| 1D9F | 605 | 53.8 | 40.52 | 14.58 | 0.58% | COMPLEX DNA POLYMERASE I KLENOW FRAGMENT - e. coli |
| 1KLN | 605 | 53.81 | 40.26 | 14.45 | 0.61% | DNA POLYMERASE I KLENOW FRAGMENT MUTANT - e. coli |
| 1NK9 | 580 | 53.39 | 40.65 | 15.35 | 0.62% | DNA POLYMERASE I - Bacillus stearothermophilus |
| 2HVH | 580 | 53.38 | 40.13 | 16 | 0.68% | in the polymerase active site Bacillus stearothermophilus |
| 2PYL | 575 | 53.85 | 38.32 | 16.39 | 0.89% | DNA polymerase complexed primer Bacteriophage phi-29 |
| 5KTQ | 543 | 54.2 | 39.61 | 15.23 | 0.64% | LARGE FRAGMENT TAQ DNA POLYMERASE Thermus aquaticus |
| 1JXE | 540 | 54.33 | 39.35 | 15.23 | 0.66% | STOFFEL FRAGMENT TAQ DNA POLYMERASE I Thermus aquaticus |
| 1QSS | 539 | 54.33 | 39.1 | 15.23 | 0.69% | LARGE FRAGMENT DNA POLYMERASE I - THERMUS AQUATICUS |
| 2OH2 | 508 | 53.22 | 37.03 | 12.77 | 1.30% | Ternary Complex of Human DNA Polymerase |
| 2R8J | 554 | 52.73 | 37.42 | 13.29 | 1.33% | DNA Polymerase eta DNA Saccharomyces cerevisiae |
| 2CW7 | 537 | 51.8 | 35.74 | 13.55 | 2.30% | endonuclease DNA polymerase thermo T. Kodakaraensis |
| 1JIH | 513 | 50.43 | 35.61 | 13.42 | 3.17% | DNA Polymerase ETA Catalytic Saccharomyces cerevisiae |
| 1G5H | 454 | 49.31 | 33.94 | 9.81 | 5.91% | murine subunit mitochondrial polymerase gamma mouse |
| 2G4C | 474 | 49.06 | 32.77 | 9.81 | 8.12% | human DNA polymerase gamma accessory subunit |
| 1V4A | 440 | 47.73 | 33.29 | 11.74 | 9.75% | N-term E. coli Glutamine Synthetase adenylyltransferase |
| 1RW3 | 455 | 48.19 | 32.65 | 11.23 | 10.19% | reverse transcriptase from Moloney murine leukemia virus |
| 3C6T | 443 | 47.47 | 33.16 | 12.52 | 10.65% | 2nd generation HIV-1 non-nucleoside reverse transcriptase |
| 1KLM | 440 | 47.22 | 33.16 | 12.65 | 11.29% | HIV-1 REVERSE TRANSCRIPTASE COMPLEXED |
| 1T94 | 459 | 47.34 | 33.03 | 10.45 | 11.29% | catalytic core of human DNA polymerase kappa |
| 2ZE2 | 428 | 46.08 | 32.52 | 12.39 | 16.84% | L100I/K103N mutant HIV-1 reverse transcriptase |
| 1TVR | 427 | 45.95 | 32.52 | 12.39 | 17.31% | wild-type HIV-1 RT+Tyr181Cys HIV-1 RT drug-resistant |
| 1BQN | 430 | 46.26 | 32.13 | 12.39 | 17.60% | mutant+wild-type HIV-1 reverse transcriptase with inhibitor |
| 1QE1 | 427 | 45.95 | 32.39 | 12.26 | 17.81% | resistant M184I MUTANT HIV-1 reverse transcriptase |
| 1UWB | 427 | 46 | 32.13 | 11.87 | 18.65% | wild-type HIV-1 RT+Tyr181Cys HIV-1 drug-resistant mutant |

**Table 5.**
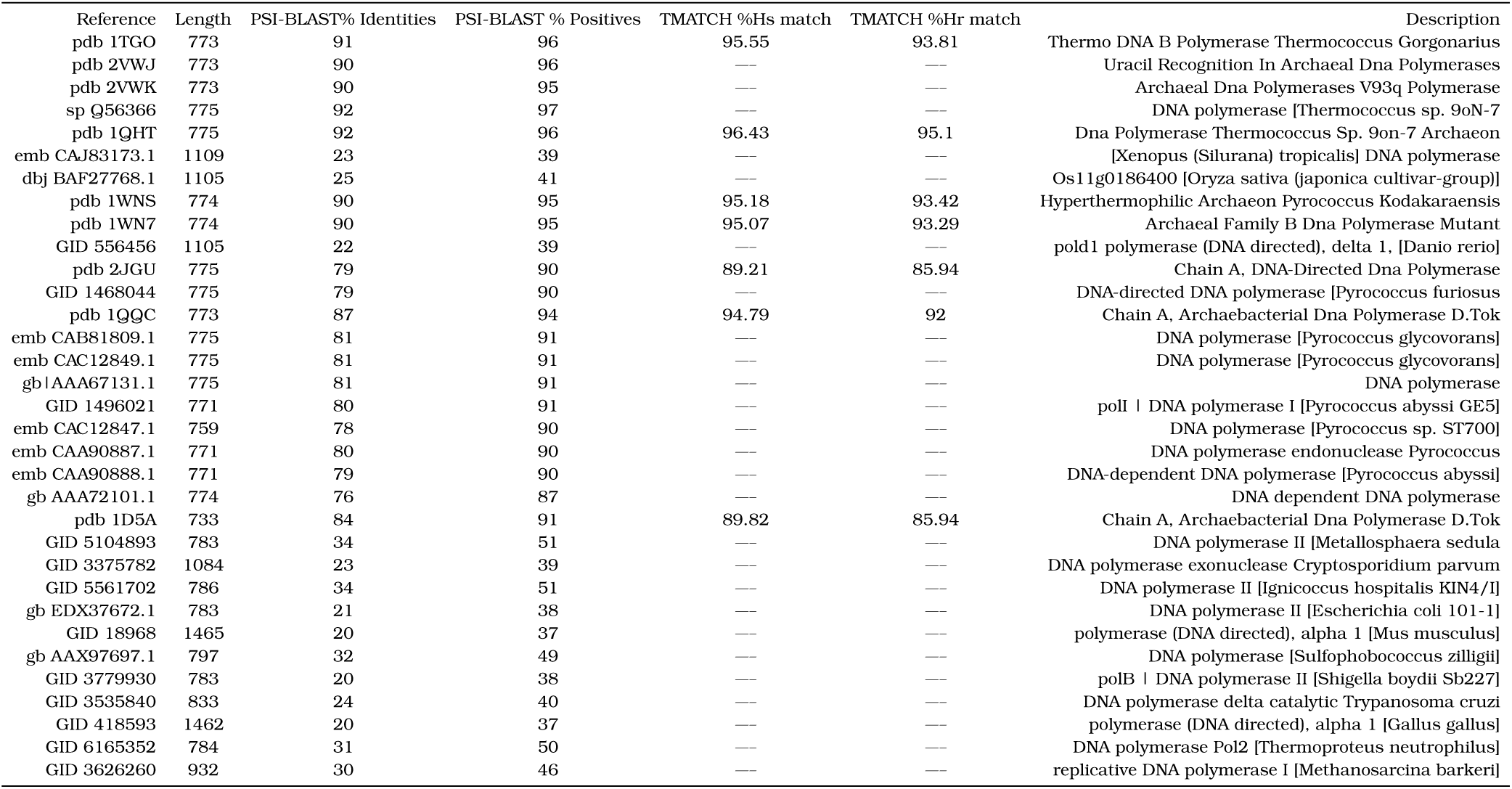
PSI BLAST of hyper-thermophilic archaeon T. thioreducens DNA polymerase B (5 iterations) with seven matches from the TMATCH table 4 for reference.Note the close values of the TMATCH % Hs and the PSI-BLAST % Positives.

In Table 3, we show results from two alignments using the the search sequence FA7_RABBIT (444 residues) and with TRYP_BOVIN (229 residues). Table 3 is sorted in ascending order by the TRYP_BOVIN sequance search alignment significance function value. The two sequences (PRTA_STRGR and PRTB_STRGR) with the highest BLAST score, lowest % identity and highest E() values, are proteases from a prokaryotic origin, thus, representing an especially challenging problem for conventional database search algorithms to recover on the first pass, yet they fall nicely within the TMATCH statistically significant alignment results. When PRLA_LYSEN is chosen as the BLAST search sequence (Pearson 1997), a number of very significant matches are obtained which exist in the BLAST TRYP_BOVIN search list from the statistically significant portion (E()< 0.2) of the list in Table 3.

The two sequences at the end of the statistically significant portion of the BLAST E() values (PRTZ_BOVIN and CFAB_MOUSE) are a little below the 25% twilight zone threshold in table 3, yet are recognized by the BLAST algorithm as being homologous to the search sequence owing to local regions of high similarity (Pearson 1997) as the BLAST statistical expectation function is designed to detect these local regions of high conservation (Altschul 1997, Karlin 1990). The PRTZ_BOVIN protein sequence result is strongly significant with both the TRYP_BOVIN and FA7_RABBIT search sequences. The CFAB_MOUSE proteein sequence result is weakly related under the TRYP_BOVIN alignment but is strongly related under the FA7_RABBIT alignment. The mixed strength of the TMATCH search significance and the marginally significant BLAST search significance for the PRTZ_BOVIN and CFAB_MOUSE illustrate the effect of size/sequence mismatch that makes members of this protein super-family difficult to recover with one alignment search As can be seen by the non-statistically significant (under BLAST) sequence KCR8_YEAST at 19.2% identity (below the twilight zone threshold), but significant under both the TMATCH search sequence alignments, the % identity alone is not a sufficient determinate of protein inter-relationships.

The first 4 sequences in Table 3 are not significant in the FA7_RABBIT sequence alignments. In table 3 columns two through five represent the TRYP_BOVIN search alignment results, with the exception that the second statistical significance, column six, represents FA7_RABBIT search sequence alignments. We repeatedly show in this section that the best choices of search sequences within a protein super-family can be difficult within some super-families, as there is a great diversity in the primary sequence, length mismatches and there is the problem of the fast roll off of percent matches within statistical transitivity relationships. In this data set the sequence lengths range upwards from a little over 1/2 the length of the FA7_RABBIT sequence, so this is part of the explanation for the differences in the two search sequence match results. However, this is not the whole story as TRY1_ANOGA is similar in length to TRYP_BOVIN and is statistically significant in the FA7_RABBIT alignments, whereas TRYP_BOVIN is not significant in the FA7_RABBIT alignments. The rest of the answer lies in the actual sequences/sequence structures themselves, the relative differences in the % identity and the sequence alignment statistical significances with different search sequences.

There is a monotonic, non-linear, relationship between the average %H (hydrophobicity) match and the f(Zb) significance, which means that large differences in f(Zb) values reflect relatively smaller, but significant differences in the %H and %I (identity) values. So the explanation of the patterns observed in Table 3 have to do with the length of the sequences, the specific sequence content, the hierarchical divergence (statistical transitivity chain) patterns within a protein family / super-family and the differing %H/ %I of each sequence relative to other sequences, keeping in mind that the %H/%I statistics are monotonically increasing metrics, but are not euclidean (e.g. orthogonal) metrics.

As of the 8th sequence, TRY1_ANOGA, the majority of the related sequences found to be statistically significant under the TRYP_BOVIN alignment also are statistically significant under the FA7_RABBIT alignments (e.g. only 5 exceptions with the TRYP_BOVIN search alignments, 10 total with the FA7_RABBIT alignments). The TMATCH FA7_RABBIT sequence alignments hold that the LORI_MOUSE is statistically weakly significant, unlike the TRYP_BOVIN alignments. The G156_PARPR sequences is not statistically significant in the FA7 RABBIT alignments, which is the same finding as with the TMATCH TRYP_BOVIN sequence alignment, and the Smith-Waterman alignment (Pearson 1997). The PLMN_PIG sequence is statistically significant under the FA7_RABBIT alignments, but not under the TRYP_BOVIN alignments.

Differences in the significance of alignment statistics can be traced to the behavior of the %Hs statistic, which is sensitive to both the number of hydrophobic proclivity fuzzy matches, and to significant mismatches in the length of the search sequence and the sequence under alignment comparison. For example, there is not a significant sequence length difference between the FA7_RABBIT sequence and the PLMN_PIG sequence. The LORI_MOUSE sequence represents a special problem within the BLAST / Smith-Waterman alignment using The TRYP_BOVIN search sequence, as the LORI_MOUSE sequence is found to be marginally related under statistical significance (E()=0.25). In this particular case, LORI_MOUSE is a pathological sequence (e.g. simple and non-homologous), containing repeats of glycine and serine of comparable size to the FA7_RABBIT sequence (Pearson 1997). With TMATCH, the LORI_MOUSE sequence is not related with TRYP_BOVIN as the search sequence, and is weakly related by statistical significance with FA7_RABBIT, the latter result being similar to the BLAST TRYP_BOVIN results. The PRLA_LYSEN sequence is not found to be statistically related to TRYP_BOVIN in the BLAST TRYP_BOVIN search (Pearson 1997), whereas it is found to be strongly related under both the TRYP_BOVIN search sequence and the FA7_RABBIT search sequence TMATCH alignments. However, PRLA_LYSEN is recovered in BLAST searches with certain other members of the Tryptophan like serine protease super-family (Pearson 1997). The advantage with TMATCH is that the number of searches with different search sequences can be reduced (sometimes very significantly) compared to other methods, as the hydrophobic proclivity scale associated with TMATCH contains more information than other search algorithms can access and the TMATCH algorithm is a global alignment algorithm, thereby accessing more information than algorithms based upon high scoring segment pairs, yet TMATCH still leverages information from the existence of high scoring segment pairs within the protein set being aligned.

The hierarchical patterns of statistically significant relationships within a protein family / super-family seen herein with TMATCH can also be seen with other algorithms such as BLAST, FASTA and Smith-Waterman. The heirarchical statistical transitivity patterns are a real phenomenon and pose significant problems for the other algorithms necessitating the need for multi-pass iterative searches. The TMATCH algorithm on the other hand is set up to use the extra information in the hydrophobicity scale to significantly cut down on the search passes needed and in many cases can be cut down to one pass. In the table 3 select sequence comparisons, we see an illustration of how a diverse protein family may need more than one search pass owing to anomalous local sequences, or anomalous results occur with pathological sequences, abnormal amino-acid frequencies and significant mismatches in protein lengths. TMATCH is likely to recover protein family members in less search passes than BLAST or FASTA because of the extra information inherent in the hydrophobicity proclivity scale.

### 3.2 A Hyper-Thermophilic Archaeon, *Thermococcus thioreducens*, DNA Polymerase B

For our next case study, we examined a Hyper-Thermophilic Archaeon (*Thermococcus thioreducens*) DNA Polymerase B using a data set of 469 sequences in FASTA format obtained from the PDB obtained by a simple text search with the character string “DNA Polymerase.” The length of these sequences ranged from 11 residues to 1220 residues long. The search sequence was a DNA Polmerase B protein from *T. thioreducens* which had 775 amino acids and was extracted from biological material recovered from the black smoker chimney at Mid-Atlantic ridge Rainbow hydrothermal vent. *T. thioreducens* is a sulfur-reducing, organo-heterotrophic strain OGL-20PT (Marsic 2008, Ng 2008, Pikuta 2007). This protein is a DNA polymerase B and the cognate DNA clamp protein will be discussed in section 3.4. (Marsic 2008, Ng 2008, Pikuta 2007)

Because of the wide variability in DNA polymerase B sequence lengths, the fact the structures of this protein family are well represented in the PDB and the fact that this particular protein function spans multiple life domains, this data set poses an excellent test of the TMATCH algorithm and the significance function f(Zb) (Marsic 2008, Dionne 2003, Imamura 2007, Nishida 2006)

The alignment statistics were sorted in increasing value (significance or lack thereof) of f(Zb) in Table 4.

Forty two of the first 73 sequences (f(Zb)<=0.181), representing 15.6% of the sequences were harvested for reproduction in Table 4. The f(Zb) scores of the aligned 469 sequences go up smoothly from 1e-7 to 1, but there are a few pathological f(Zb) values outside of the interval of 0 to 1, where both %Hr and %Hs are very low (both <4%). It is very interesting to note that the rank percentage at 15.6% was very close to the f(Zb) significance score at that point with f(Zb)=18.1%.

The value of the function f(Zb) for the 1st seven sequences of the data set is very highly significant at 1.0e-7 to 3.6e-7, with the %Hs and %Hr both greater than 85%. After the first 7 sequences the %Hr and %Hs values drop significantly to 50-60%. The first 7 sequences are all thermophilic archaeons. The percent identity of these 7 sequences begins about 4% lower than %Hr, and then declines to about 10% lower than the %Hr statistic. After the first 7 sequences, the %identity immediately declines to be typically within 2-6% of the square of the %Hr statistic. During the 1st seven sequences the %Hs and %Hr statistics are within 1-4% of each other.

This data set has a number of sequences with %identities transitioning from well above to below the twilight zone. While the percent identity is at 20%, the hydrophobicity based match statistics %Hs and %Hr are running around 50%, demonstrating the significant information content of the hydrophobicity scales, which readily detects the substitution of similar amino-acids. When we examine the performance of the statistical significance function sequences of relatively low percent identity with the search sequence, we notice that the relationship with the search sequence is quite significant and *would not be found as such with the statistical significance function used by BLAST and FASTA because of the low sequence identities*.

Starting at the 8th sequence, the %Hr statistic steadily goes from around 7% less than the %Hs statistic to to around 16% less than the %Hs statistic at the f(Zb)=10 %. Up to this point, it is primarily the decline of the more conservative %Hr statistic that drives the decline in the f(Zb) significance (e.g. higher value). After f(Zb)=10 %, the %Hr statistic stabilizes and runs around 14 - 16% less than the %Hs statistic, for an overall average difference of ∼14%. By the time the %identity declines to around 11.2% (%Hr about 32.7%), the f(Zb) statistic goes above 10 % and is deemed to be statistically insignificant. An inspection of the sequence descriptions shows that the actual transition to non-relatedness begins at f(Zb)=8.12-9.75 % (italicized font in Table 4).

The 8th sequence in order of statistical significance by f(Zb) (at 2.2e-4) is an archaeon, though not a thermophile.*The 9th sequence in order of statistical significance by f(Zb) (at 4*.*6e-4) is a surprise, which is an HIV type 1 DNA polymerase*. The nature of this surprise is that there is a relationship between the DNA replication machinery between a hyper-thermophile archaeon and a human virus, which would imply that either the genesis of this virus dates back very early in evolutionary time or that we see a type of convergent evolution. The former explanation seems the most plausible as we have thermophilic bacteriophages represented in the alignments strongly allied with the *T. thioreducens*.

From the 9th sequence up though where f(Zb) < 2.3% the sequences are mostly associated with bacterial DNA polymerases, some of them thermophiles, but a few sequences belong to bacteriophages. Right at f(Zb)= 3.2 % there is a DNA polymerase corresponding to a yeast *Saccharomyces cerevisiae*.

Between f(Zb)> 10.2 % and f(Zb)< 18.7 %, there are a number of reverse transcriptases, representing a couple of viruses (including HIV), the mouse mitocondrial and human DNA Polymerase gamma / kappa. The reverse transcriptases, are in the significance range being ascribed to being distantly related to probably not related, which considering their structure and function is true in this case. There are a number of sequences ostensibly in the non-significant region of f(Zb) above 10 % and less than 30 %, which are reverse transcriptases, which still have some resemblance to the search sequence structure, but not as closely as the statistically significantly related sequences. The structure of the hyper-thermophile search sequence was compared to all of the sequence structures in Table 4, and they all shared a common architecture (alpha + beta) and basic scaffold (core fold morphology). However, there was increasing divergence in the structural detail noted, which tracked the f(Zb) significance scores.

### 3.3 A hyper-thermophilic archaeon, *T. thioreducens*, PCNA DNA Clamp and Bacterial DNA Clamps

Cellular life from the life domains of Archaea, Procarya and Eucarya possess sliding clamps used to tether their replicative DNA polymerases to single stranded DNA during cellular replication. In bacteria this function is accomplished through the beta clamp and the analogous function in Archaea and Eucarya is performed with the Proliferating Cell Nuclear Antigen (PCNA). PCNA is also used as a scaffold for ancillary processing factors like DNA ligase I and the flap endonuclease and as an organizing factor for processing factors used to rescue stalled strand copying. These several processing factors, including the DNA polymerase that binds to the inter-domain connector loop on the PCNA protein. There is limited sequence / super-secondary, domain structural homology between the bacterial beta clamps and the Archaea / Eucarya PCNA proteins, however, there is a strong quaternary structural homology between the beta clamp and PCNA oligomers. The quaternary structure of these two protein families are strongly similar, having a toroidal morphology with quasi-six fold symmetry possessing a central hole through which the single DNA strand moves. The bacterial beta clamp, sometimes known as the DNA Polymerase III beta sub-unit, is a dimeric (three domains per monomer) ring, while the Achaea / Procarya PCNA proteins is a trimeric (two domains per monomer) ring. (Dionne 2003, Imamura 2007, Nishida 2006) We studied two PCNA protein sequences from two species belonging to Domain: Archaea Group: Euryarchaeota, but are from different thermal extremes (Byrne-Steele 2008). The psychrophilic / psychro-tolerant (cold loving) archaeon *Methanoccoides burtonii* (Mburt) has a growth range of 1-28°C, with an optimal temperature of 23°C. The hyper-thermophilic archaeon *Thermococcus thioreducens* (Tthio) has an optimal growth temperature range from 82-85°C. For comparison purposes with the archaon PCNA proteins, the bacterium *Shigella boydii* (Sboyd) beta clamp protein is used. The two archaeon PCNA sequences were BLAST searched against the NCBI protein database, with the Tthio sequence returning no hits, but the Mburt sequence returns one hit with the Sboyd sequence lcl7426 unnamed protein product, length=245. (Byrne 2008, Marsic 2008; Ng 2008, Pikuta 2007)

The TMATCH results are also seen below, with the best match being between the Mburt and Tthio PCNAs (signicance = 9e-05) versus the Mburt PCNA to Sboyd beta Clamp match being signicant (5.9e-04), but not as good as the two archaeon PCNA alignments as would be expected. The BLAST alignment significance (E() = 0.018) is the most significant of the 3 HSSP’s retrieved by BLAST, yet it is not nearly as significant (by orders of magnitude) as the TMATCH alignment. Both algorithms (TMATCH and BLAST) would have found the Sboyd Clamp protein, but the BLAST results would have looked more like a suggestion than a positive hit as found by TMATCH. The BLAST alignment / match results of Mburt vs. Sboyd are three HSSP’s with statistical expectation / significance values of E1()=0.018, E2()=0.088 and E3()=0.47 respectively. BLAST suggests HSSP \ #1 is between the middle of the Mburt PCNA and into the middle of the Sboyd beta clamp. BLAST also suggests that HSSP \ #2 lies in the middle of the Mburt PCNA protein sequence just ahead of the HSSP \ #1 segment and corresponds to a segment in the latter part of the Sboyd beta clamp, which is a curious type of location inversion. The third HSSP is of marginal relevance.

### 3.4 NCBI BLAST results for Mburt vs. Sboyd

These are the BLAST alignment results for the three HSSP regions of the BLAST search alignment. Score = 21.9 bits (45), Expect = 0.018, Method: Compositional matrix adjust. Identities = 19/66 (28 %), Positives = 29/66 (43 %), Gaps = 7/66 (10 %)

~~~
Query 220 SNNIRAHVGDFIFTSKLVDGRFPDYRRVLPKNPDKHLEAGCDL---- 263
          S++ +GF +T L+D P R P+ P L A L
Sbjct 94 SKKLKIQIGGFSYTISLLD---PSTIRAEPRIPQLELPAEIVLNGKD 138
-----------------------

Query 264 LKQAFARAAILSNLKQAFARAAILSNEKFRGV 285
          L++A A+ S+L++A A +S+ GV
Sbjct 139 LQKAVKAAEKISDLQKAVKAAEKISDHMLLGV 159
-----------------------
~~~

Score = 19.6 bits (39), Expect = 0.088, Method: Compositional matrix adjust. Identities = 10/42 (23%), Positives = 24/42 (57%), Gaps = 1/42 (2%)

~~~
Query 310 YSGAEMEIGFNVSYVLDVLN-ALKCENVRMMLTDSVSSVQIE 350
          + + E+G ++S + D+L A + + V+M L + ++I+
Sbjct 59 FEANDCELGLDLSRINDILGVADRDDKVQMELDEESKKLKIQ 100
-----------------------
~~~

Score = 17.3 bits (33), Expect = 0.47, Method: Compositional matrix adjust. Identities = 5/20 (25%), Positives = 10/20 (50%), Gaps = 0/20 (0%)

~~~
Query 209 GGDNPLRVQIGSNNIRAHVG 228
          G D P+++ N + +G
Sbjct 216 GNDFPIKINFSIANGKGTIG 235
~~~

The BLAST search results effectively have three high scoring segment pairs of moderate percent identity at 23-28%, although the “positives” add similar property amino-acids to the total for a higher percentage at 43-57%. We see that the BLAST algorithm for positives detection does to some extent capture the similarity scale reflected by the hydrophobicity proclivity metric.

### 3.5 TMATCH alignment results

Alignment search (row) sequence: Mburt PCNA *Methanoccoides burtonii* (psychrophilic) cold archaeon The column string: TthioPCNA *Thermococcus thioreducens* hyperthermophilic archaeon

Output of the aligned row/column string:

~~~
MburtPCNA> M-FKATIDAY-LL-K--DSIETLSVLVDE-ARFRISPEGVV-VRAVD
TthioPCNA> MPFEIVFDGAKDFADLIAT-AS-NLI-DEAA-FKITEEG-ISMRAMD
MburtPCNA> -PANVAMVSFDLT-PE-AF-D-DFEANDCE-LG--LDLSRINDILGV
TthioPCNA> PS-RV-VL-IDLNLPESIFSKYEVE-EE-ETIGINMD-H-FKKILKR
MburtPCNA> ADRDD-KVQMELD-EESKKLKIQIG--GFSY--T--ISLLDPSTIRA
TthioPCNA> GKNKDTLI-LR-KGDE-N-F-LEVTFEG-TAKRTFKLPLIE---VE-
MburtPCNA> E-P-RIPQLELPAE-IVLNGKD-LQK AV-KA--AEK-ISDHMLLGV
TthioPCNA> ELELDLPELPFTAKVVVL-G-EVLKE-AVKDASLVSDAL-K-FIAT-
MburtPCNA> EGESFFMEAEGDTDRVKLTMT-RDQ-LIDIK-PSQVRS--LFS-LDY
TthioPCNA> ENE-FTMKAEGETNEVEIKLTLEDEGLLDLEVEEETKSAYGISYL-A
MburtPCNA> LSDIIKPA-SKSNE ISLHLGNDFPIKINFSI-ANGKGTIGYL
TthioPCNA> --DMIK-GIGKADEVI-IRFGNEMPLQMEYPIRDEGK-LI-FL
MburtPCNA> LAPRIESD
TthioPCNA> LAPRVE-D
~~~

Number of gaps=97, % gaps=26.57, Avg score fitted % Hst match=68.0221%, % Hs match =66.08392, % Hr =57.8947, **Significance =5.8861e-05**

~~~
------------------
~~~

Alignment search (row) sequence: MburtPCNA *Methanoc-coides burtonii* psychrophilic cold archaeon The column string: SboydCLMP *Shigella boydii* bacteria

Output of the aligned row/column string:

~~~
MburtPCNA> M-F---K-A---TI-D--AYLL-KDS-IET-L-SVLVD--E-ARFR
SboydCLMP> MKFTVEREHLLKPLQQVSG-PLGGRPTL-PILGNLLLQVADGT-LS
MburtPCNA> I-SPE-GV-VV-R-A-V---DP-A-NV-A--MV-SFD-LTPE-A-F
SboydCLMP> LTGTDLEMEMVARVALVQPHEPGATTVPARKFFDICRGL-PEGAEI
MburtPCNA> -DDFEANDCELGLDLSRINDI-L-GV--AD-RD-DKVQMELD---E
SboydCLMP> AVQLEGERM-L-VRSGR-SRFSLSTLPAADFPNLDDWQSEVEFTLP
MburtPCNA> E-S-KKLKIQIGG-FS-YT------I-S-LLDPS-T--IR--A-E-
SboydCLMP> QATMKRL-IE-ATQFSMAHQDVRYYLNGMLF-ETEGEELRTVATDG
MburtPCNA> PR--I-P-QL-E-L-PAEIVLNGKD-LQK-AVKAAEKISD-HM-L-
SboydCLMP> HRLAVCSMPIGQSLPSHSVIV-PRKGVIEL-MR-MLDGGDNPLRVQ
MburtPCNA> LGVEGESFFMEAEGDTD-RV--K-L-T-MTR-DQLI-DIKP-SQVR
SboydCLMP> IG-SNN--I-RA-HVGDFIFTSKLVDGRFPDYRRVLPK-NPDKHLE
MburtPCNA> -S--LF-S-L-D-Y-L-SD-I--IKPA-SKSN-E -ISLHLG-N-D
SboydCLMP> AGCDLLKQAFARAAILSNEKFRGVRLYVSENQLKITAN-N-PEQEE
MburtPCNA> -FPIKINFSIA-NG--K-G-TIGY-L--L-A---PRIE---SD---
SboydCLMP> AEEI-LDVT-YSGAEMEIGFNVSYVLDVLNALKCENVRMMLTDSVS
MburtPCNA> -----------------------------
SboydCLMP> SVQIEDAASQSAAYVVMPMRL
~~~

Number of gaps=183, % gaps=42.42%, Avg score fitted % Hst match =66.29%, % Hs match =55.59%, % Hr =57.89%, **Significance =3.5750e-04**

~~~
-----------------------------
~~~

It can be seen from the TMATCH alignment results above of the two PCNA DNA clamps that the PCNA *Methanoccoides burtonii* (psychrophilic) cold archaeon has an excellent alignment with the *Thermococcus thioreducens* hyperthermophilic archaeon, which is not unexpected. However, what is a little bit more of a suprise is how well that the *Methanoccoides burtonii* PCNA aligns with the *Shigella boydii* eubacteria beta clamp. This latter alignment illustrates how adept that the TMATCH algorithm with the hydrophobicity proclivity scale is at finding the high alignment similarity at both the local and global levels.

### 3.6 PSI-BLAST Results of hyper-thermophilic archaeon *T. thioreducens* DNA polymerase B

PSI-BLAST is one of the most commonly used algorithms for finding distant protein relationships. PSI-BLAST detects distant relationships with an alignment algorithm possessing a strong ability to recover remote structural (common fold) homologues. The PSI-BLAST algorithm uses an iterative procedure to improve the detection of distantly related proteins. At the end of each iteration, PSI-BLAST does a multiple alignment of the protein sequences recovered during that iteration against the database. PSI-BLAST then produces a Position Specific Scoring Matrix (PSSM) from the multiple alignment of sequences. The PSSM is a pattern or template used for the next iterative search of the database for additional similar sequences. The PSI-BLAST iterations proceed until the protein search converges (e.g. no new proteins recovered) or the maximum number of iterations stipulated by the user is reached. With an adequate number of iterations, weak but biologically meaningful similarities may be detected. Obviously, the accuracy of the local and multiple alignments are crucial to detecting distant relationships where sequence identity could fall as low as 10-12%. Recent studies of the PDB have reported that around 75% of newly determined structures with no protein matches in the PDB fall into already known fold families. (Altschul 1997, Altschul 2005, Friedberg 2000)

Multiple-alignment / pattern generating algorithms have been demonstrated to recognize three times as many remote homologs as conventional pairwise methods. In one study, PSI-BLAST alignment performance and accuracy were characterized against databases of carefully selected and aligned (structural and sequence alignments) distantly related proteins on the basis of sequence, but sharing a common fold. The accuracy of the PSI-BLAST alignments were calculated as the percentage of positions correctly aligned in both the reference sequence and structural alignments, with the accuracy being 43.5 ± 2.2% for the first run and improving to 50.9 ± 2.5% within five iterations. (Friedberg 2000)

There are seven PDB matches taken from the TMATCH DNA polymerase B proteins in Table 4 that were harvested from the Protein Data Bank and selected for comparison with a PSI-BLAST search, which is represented in table 5. The PSI-BLAST % Positives reflect a match statistic formulated from both identical and highly similar amino-acids.

The PSI-BLAST % Identities and % Positives bound above and below the TMATCH hydrophobicity match statistics %Hs and the more conservative %Hr statistic, illustrating the reasonableness of the hydrophobicity scale. The % Identity (not shown in Table 5) between the two methods is either the same or 1% different. At 75% Identity and above, the square of the PSI-BLAST % Positives (the estimated % identity column) is very close to the % Identity. The square of the PSI-BLAST % Positives are still reasonably close to the % Identities, even as low as 30% Identities. An inspection of the Table 5 shows that there are a number of the highest scoring matches which have low sequence identities (20-25%), illustrating the PSI-BLAST focus on local regions of similarity, recovering individual domains and secondary / super-secondary structure. On the other hand TMATCH represents a global alignment algorithm, which is also is sensitive to structure that results from domains and secondary / super-secondary structure forming the diagonals in the match matrix reflecting sequences of high similarity.

## 4 DISCUSSION

In Table 6 (from Cavanaugh 2009C), the relationships between %Hs and %Hr may be observed for sequences of lengths closer than with the Tryptophan like Serine Protease super-family examined above. The Table 6 statistics are harvested from experimental runs (data not shown) completed and compiled for this discussion. The first thing to be noticed is the reasonably close relationship between the %Hst, %Hs and %Hr statistics. Notice that the binomial (approximation) f(Zb) cumulative probability distributions do not roll off as fast as does the E() expectation / significance function (last column of Table 6) typically used for database search alignment algorithms, like BLAST and FASTA, so the results of the binomial distribution must be interpreted somewhat differently than the E() function, as is reflected in the f(Zb) significance relation. Since the Hydrophobicity proclivity scale reflects the physico-chemical similarity of amino-acids, it is reasonably to be expected that there is less of a problem with ambiguity of sequence statistical relationships around the twilight zone threshold (25% identity).

**Table 6.**
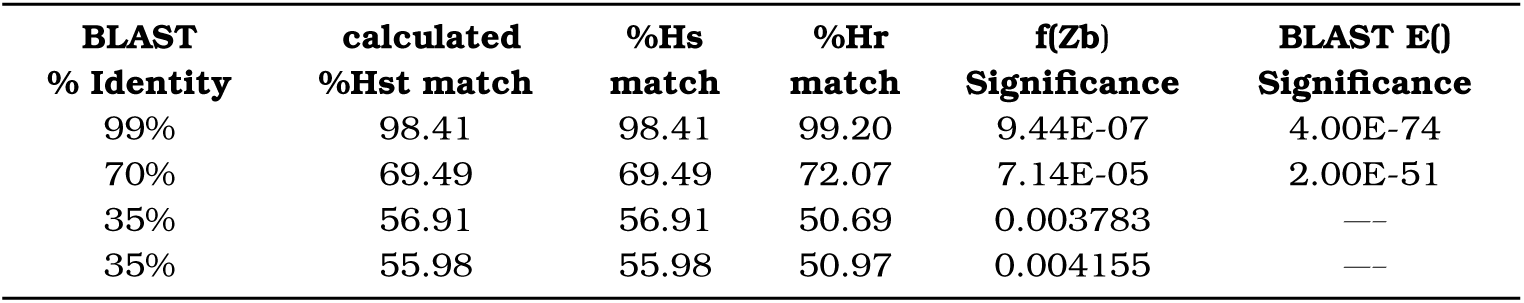
Comparing TMATCH statistics versus BLAST statistics at BLAST 99% Identity, 70% Identity and 35% Identity.

As can be seen in table 6, the 35% identity sequences are really more closely related than would be apparent because of the physico-chemical similarities reflected in the %Hs and %Hr metrics, but are not highly reflected in the BLAST % Identity (Cavanaugh 2008b, Cavanaugh 2008c). As can be seen in the PSI-BLAST vs. TMATCH results in Table 5, TMATCH compares very favorably with the results of the PSI-BLAST algorithm in terms of effectiveness. TMATCH has other properties that give it an edge over PSI-BLAST, such as being a global alignment algorithm as well as being sensitive to local areas of similarity between two proteins being aligned. TMATCH has a speed advantage for protein database searches by using Sa to estimate %Hst, therefore not having to extract the alignment with attendant over-head and possessing the ability to cut down the number of iterations through large protein databases in order to find distantly related proteins.

Additional work (not shown, Cavanaugh 2008c) benchmarked the TMATCH search global alignment algorithm against the FASTA local search alignment algorithm with three carefully curated search alignments with three different search proteins (Pearson 1995).

1. GTM1_HUMAN Glutathione S-Transferase HB Subunit 4 (P094881, UniParc, 218 AA long)
2. OPSD_HUMAN Rhodopsin family of GProtein Coupled Receptors (P081001, UniParc, 348 AA long)
3. CAR1_DICDI CAMP Receptor 1 protein (P137731, UniParc, 392 AA long)

Within the OPSD_HUMAN G protein coupled receptor protein search alignment, there were 3 different protein families of Gprotein coupled receptors. The largest family is the rhodopsin family, including cationic amine receptors like dopamine and acetylcholine, prostanoids, ordorants and a number of other hormones. The other two families are the glutamate receptor family and the secretin/calcitonin family (Pearson 1995). In the CAR1_DICDI CAMP Receptor 1 protein alignment search there were Rhodopisin family, Calcitonin family and Secretin family proteins (Pearson 1995). Generally speaking, with these two alignment searches TMATCH performed as well or better than FASTA, particularly where there are remote homolgies with low sequence percent matches. There are two possible significance functions that can be used with TMATCH, based upon either the %Hst statistic or the average of the %Hs and the %Hr statistics. When the size mismatch between the search and a comparison sequences hits about 2x, the latter statistical significance percentage based upon the average of average of %Hs and %Hr starts to become artificially depressed, even through the fuzzy match identities are still reasonably good, and may not pick up a remote homologe that would be detected with the statistical significance function based upon the %Hst statistic.

In the CAR1_DICDI CAMP Receptor 1 protein search alignment, the TMATCH algorithm generally exceeds the performance of the FASTA algorithm for detecting homologous relationships. The marginally significant FASTA search results trapped some Cytochrome B protein sequences, which were found to be statistically significant matches with TMATCH, which posses an interesting and significant disconnect between the TMATCH and FASTA results. The Cytochrome B protein is an integral membrane protein with 8 or 9 transmembrane alpha helices intercalated together (Esposti 1993, Widger 1984). The G protein coupled receptor protein is an integral membrane protein with 7 transmembrane alpha helices intercalated together (Atwood 1994, Palczewski 2000). Comparison of the fundamental/general fold of these two protein families illustrates a good example of basic protein family scaffolds, reflecting the structural and environmental factors for which the proteins must be adapted. Protein families with closely related scaffolds would be expected to have significant matches with alignment algorithms (and the related scales) that are sensitive to structure and structural homologies.

## 5 CONCLUSION

We have described a global alignment algorithm for searching proteins using a hydrophobic proclivity index that allows for the identification of proteins with similar secondary and tertiary structure. The TMATCH alignment algorithm makes several important adaptations over existing alignment algorithms based upon the classical dynamic programming algorithm, including the dropping of affine gap extension penalties.

The BLAST and FASTA algorithms are optimized to recover protein sequence relationships based upon conserved motifs reflected in local regions of high similarity. The fundamental hydrophobicity proclivity metric of amino-acid similarity leads to some profoundly different structural adaptations in the TMATCH algorithm, which was designed and constructed around such a scale. The TMATCH algorithm does share many features with the FASTA and Smith-Waterman alignment algorithms as they are all dynamic programming algorithms, but there are significant differences. The FASTA and Smith-Waterman algorithms use an affine gap penalty function, whereas in TMATCH we use a constant gap penalty. The gap extension penalty aspect of the affine gap penalty is implemented in a different fashion within TMATCH with the favorable diagonal transition reward score, and unfavorable transition punishment score, which causes the algorithm to prefer to go along diagonals instead of continuing along a row or column causing gaps, but extends gaps where necessary to pick up new similarity diagnonals.

The TMATCH algorithm shares a fundamental similarity with the FASTA algorithm, in that each algorithm starts with a dot plot style analysis. The FASTA algorithm uses diagonal clusters as sites to nucleate local alignments to recover sequence segments of high similarity. TMATCH uses the diagonal clusters as guide posts or milestones for tracing the preferred / best path through the score matrix, which is seeking to recover a best trace based upon a more global or distributed level of similarity, as opposed to simply finding pockets of local sequence similarity. Therefore, the TMATCH algorithm is leveraging the fact that amino-acids of similar physico-chemical properties tend to cluster together in real proteins. Proteins with similar segments of amino-acids with similar physico-chemical properties, strongly tend to assume similar tertiary structure, which is the most robust notion or basis for the recover protein family / super-family homologies.

An algorithm such as TMATCH and the associated hydrophobicity scale that recovers amino-acid alignment relationships across a wide swath of the two sequences being aligned, should be expected to reflect secondary structure and super-secondary structure affinities (Cavanaugh 2008a, Cavanaugh 2008c, Pintar 2003a, Pintar 2003b). In many protein families, the relationship is probably expressed as the result of convergent evolutionary adaptation or derivation from an ancient homolog found in similar structural circumstances. Alignment based searches are often used to recover homologous relationships for an unknown protein sequence in order to understand function and/or structure. Common scaffold (e.g. have a common fold structure but are not homologous) alignment confounding could occasionally be a problem when trying to uncover functional relationships, but this seems a small price to pay for the ability to reliably recover structural relationships. If an alignment can find a known structure which possesses common scaffold similarity to the structure of the unknown protein, then it may still serve as an excellent starting point to understand the unknown protein cellular environment or cellular location, and may serve as a starting point to begin to refine the structure of the unknown protein.

## ACKNOWLEDGEMENT

*Funding:*

